# Single-shot phase contrast microscopy using polarisation-resolved differential phase contrast

**DOI:** 10.1101/2021.04.14.437846

**Authors:** R. Kalita, W. Flanagan, J. Lightley, S. Kumar, Y. Alexandrov, E. Garcia, M. Hintze, M Barkoulas, C. Dunsby, P.M.W. French

## Abstract

We present a robust, low-cost single-shot implementation of differential phase microscopy utilising a polarisation-sensitive camera to simultaneously acquire 4 images from which the phase gradients and quantitative phase image can be calculated. This polarisation-resolved differential phase contrast (pDPC) microscopy technique can be interleaved with single-shot imaging polarimetry.

## 1. Introduction

Phase microscopy is attracting significant interest with the development of quantitative or semi-quantitative techniques that can provide label-free means to image cell dynamics, to aid cell segmentation and or to provide readouts for diverse potential applications^1^ such as blood screening, cell sorting, cell tracking, microbiology, neuroscience, and pathology. An application of wide potential interest could be to provide label-free segmentation on existing fluorescence microscopes. There are many approaches to realize quantitative phase microscopy (QPM) including explicit interferometric techniques such as phase stepping interferometry, off-axis (digital) holography, shearing interferometry, techniques that reconstruct the phase of the electric field scattered by the sample from intensity measurements, such as transport of intensity^2,3^ and (Fourier) ptychography^4^, and non-interferometric wavefront sensing approaches including the use of Shack Hartman sensors^5^ or other phase gradient measurement techniques^6^, including differential phase contrast microscopy^7,8^. Most of the phase contrast techniques reported to date require specialist optical components (e.g. objective lenses with phase rings, Nomarski prisms, wavefront sensors), significant modification of the microscope configuration (e.g. for interferometry) and/or the acquisition of multiple images to calculate the phase contrast image. This can make it challenging to implement them on existing fluorescence microscopes.

Differential phase contrast (DPC) techniques can be conveniently implemented on existing microscopes by modifying the illumination path or imaging path^7^ and do not require special objective lenses. The illumination approach entails acquiring transmitted light images for different illumination or detection angles, from which semi-quantitative phase images can be reconstructed. An elegant approach to DPC phase contrast microscopy can be conveniently realised by acquiring transmitted light images using four illumination directions constrained by semicircular masks in the back focal plane of the condenser lens, as indicated in Figure 1. This can be conveniently implemented using a liquid crystal device (LCD) in the back focal plane of the condenser lens^9^. Alternatively, this oblique illumination can be provided using a programmable LED array as the illumination source^8^. The phase contrast image may be calculated from a set of four transmitted light images acquired with the condenser pupil plane masks illustrated in Figure 1. The phase gradient images serve as the input data to the convolution calculation outlined in [8], which can be conveniently undertaken using the DPC software^10^ shared by the authors of reference 10.

**Figure 1.**
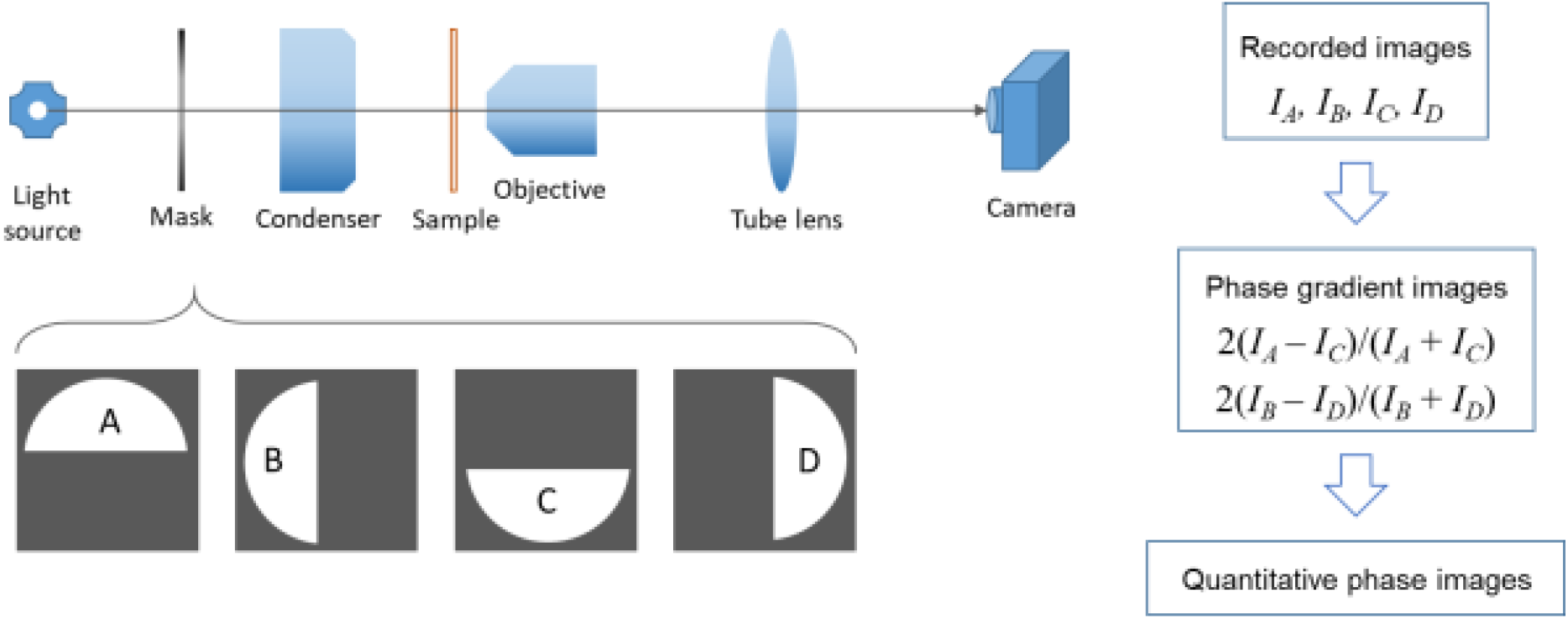
Schematic of differential phase microscopy with semicircular masks in the pupil plane of the condenser lens

Usually such DPC techniques require the sequential acquisition of two orthogonal pairs of images with semicircular apertures, each rotated by 180^0^, in the condenser lens pupil plane. We were motivated to implement a version of DPC microscopy that could acquire quantitative phase images in a single shot, rather than requiring 4 sequential image acquisitions. This would be particularly important to avoid motion artefacts, e.g. when imaging dynamics of live cells or cell sorting. A single-shot implementation DPC microscopy has been reported that utilised three colour filters and a colour camera with an RGB Bayer filter sensor^11^. This “cDPC” approach provides single-shot quantitative phase imaging but it is not suitable for spectrally resolved imaging because it is already utilising multiple spectral channels and it can be challenged by chromatic aberrations or spectral variations in sample properties. It could also be difficult to implement this in a fluorescence microscope without switching the fluorescence filter cube out of the beam path.

We reasoned that if we replaced the conventional camera with a polarisation sensitive camera using the Sony “Polarsens™” IMX250MZR sensor, we could acquire 4 images simultaneously at up to 75 frames per second without any spectral trade-offs. We refer to this technique as polarization-based differential phase contrast (“pDPC”). Figure 2 presents the principle of pDPC microscopy, which we implemented on a low-cost, modular, open microscopy platform we describe as “*openFrame*” ^12^, using a plug-in developed for Micro-Manager ^13^ that acquires and processes the polarisation-resolved image data. We aimed to obtain the orthogonal phase gradient images, 2(*I*_*A*_*-I*_*C*_)/(*I*_*A*_*+I*_*C*_) and 2(*I*_*B*_*-I*_*D*_)/(*I*_*B*_*+I*_*D*_), to serve as input data for the DPC software^10^ and to capture them in a single image acquisition using the Polarsens™ camera (BFS-U3-51S5P-C FLIR Systems Inc). Accordingly we fabricated a composite polarizer mask to be located in the condenser pupil plane by mounting 4 pieces of polarizing film (Edmund Optics, #86-179) on a glass substrate with the polarizer axis for each quadrant orientated at 0^0^, 45^0^, 90^0^ and 135^0^. For a single image acquisition, the sample is simultaneously illuminated by beams coming from 4 different directions with 4 different polarisation orientations that are aligned with the 4 polarisation directions of the pixels on the Polarsens™ camera chip. The four illumination beams contribute intensity distributions *I*_*α*_, *I*_*β*_, *I*_*γ*_, *I*_*δ*_, to the image acquired on the camera. As indicated in Figure 2(a, b), the four polarisation-resolved sub-images images recorded on the Polarsens™ camera (*I*_*1*_, *I*_*2*_, *I*_*3*_, *I*_*4*_) can be used to calculate phase gradient images, 2(*I*_*A*_*-I*_*C*_)/(*I*_*A*_*+I*_*C*_) and 2(*I*_*B*_*-I*_*D*_)/(*I*_*B*_*+I*_*D*_), from which the quantitative phase image can be calculated using the DPC software^10^, which is run in MATLAB as a subroutine called from the MicroManager plug-in. The application of pDPC is illustrated with exemplar images of HEK cells acquired with a 0.3 NA 10x microscope objective lens. As explained in reference 8, this approach does not reconstruct low spatial frequency components of the phase contrast image and so the reconstructed phase values are relative to an unknown offset.

**Figure 2.**
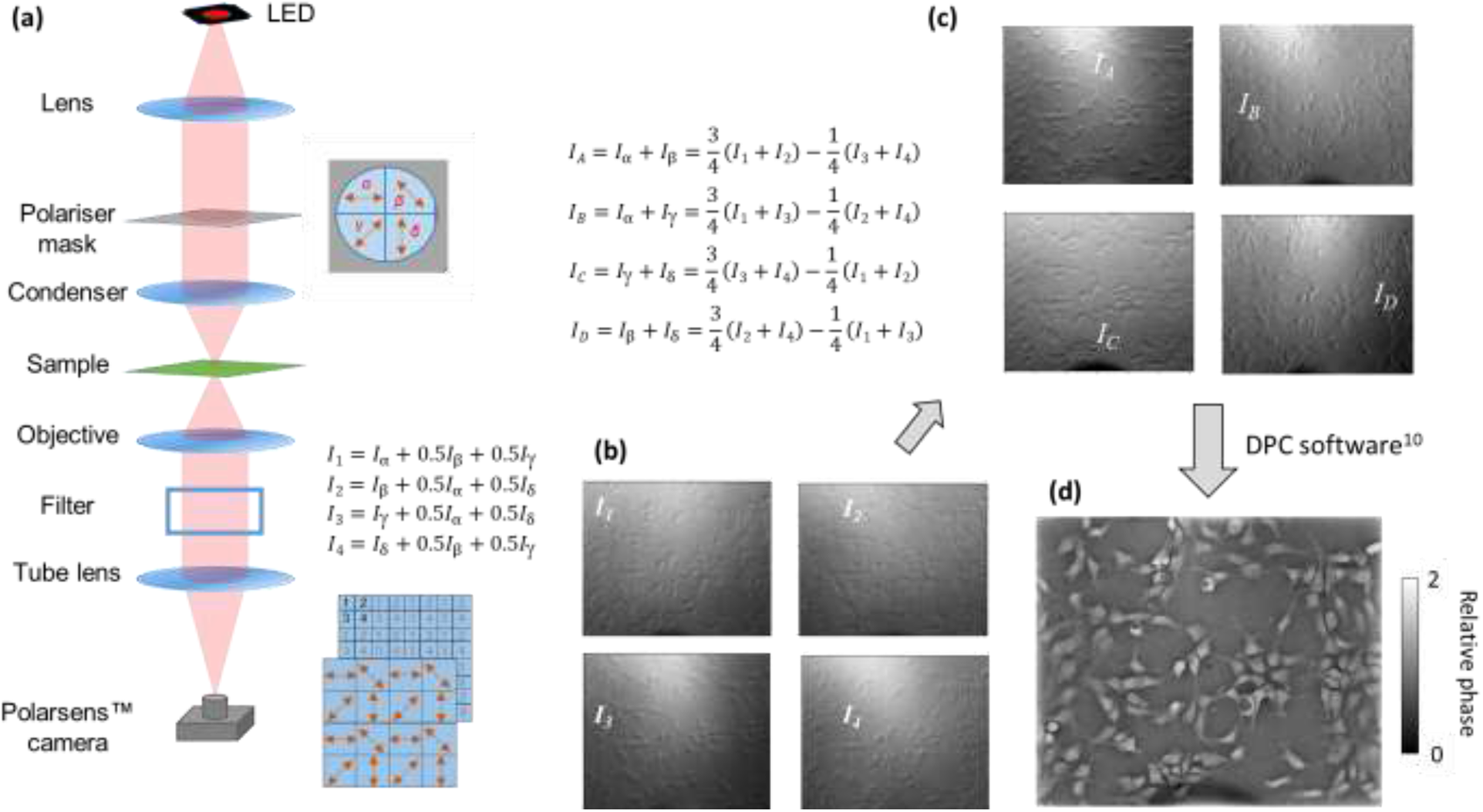
Schematic of single shot “pDPC” microscope utilising a composite polarizer in the back focal plane of the condenser lens for which the four quadrants contain polarizing film orientated at 0^0^, 45^0^, 90^0^ and 135^0^. The image acquired on the Polarsens™ camera can be divided into 4 polarisation-resolved images, I_1_, I_2_, I_3_, I_4_.

Figure 2 presents an exemplar quantitative phase image of fixed HEK cells calculated from data acquired in a “pDPC” single image acquisition of 30 ms integration time using a 0.3 NA 10x microscope objective lens. The illumination source as a white LED and the transmitted light images were recorded via a fluorescence filter cube in place to provide a detection channel centered on 590 nm ±55 nm. Figure 3 shows pDPC image data of a similar sample acquired in the same microscope but also using a 10 nm bandpass filter centred at 560 nm; two expanded views of regions of interest are shown with linear phase profiles corresponding to the yellow lines across them. This illustrates the potential for rapid semi-quantitative phase imaging, which we are exploring to monitor cell dynamics, for high content analysis and for histopathology. We note the potential for broadband operation of this approach is limited only by the properties of the polarizers, in contrast to the cDPC approach^11^ utilising an RGB sensor, or to the LCD based approach of [9] where the modulation function of an LCD modulator may be limited to a spectral range of ∼50 nm full width half maximum. Using a suitable illumination source, the broadband nature of our polarisation-based approach to DPC would enable it to be implemented on a fluorescence microscope without changing the fluorescence imaging components; this would facilitate rapid sequential or simultaneous acquisition of fluorescence and phase-contrast images on the same camera without needing rapid switching of the fluorescence filters in the beam path.

**Figure 3.**
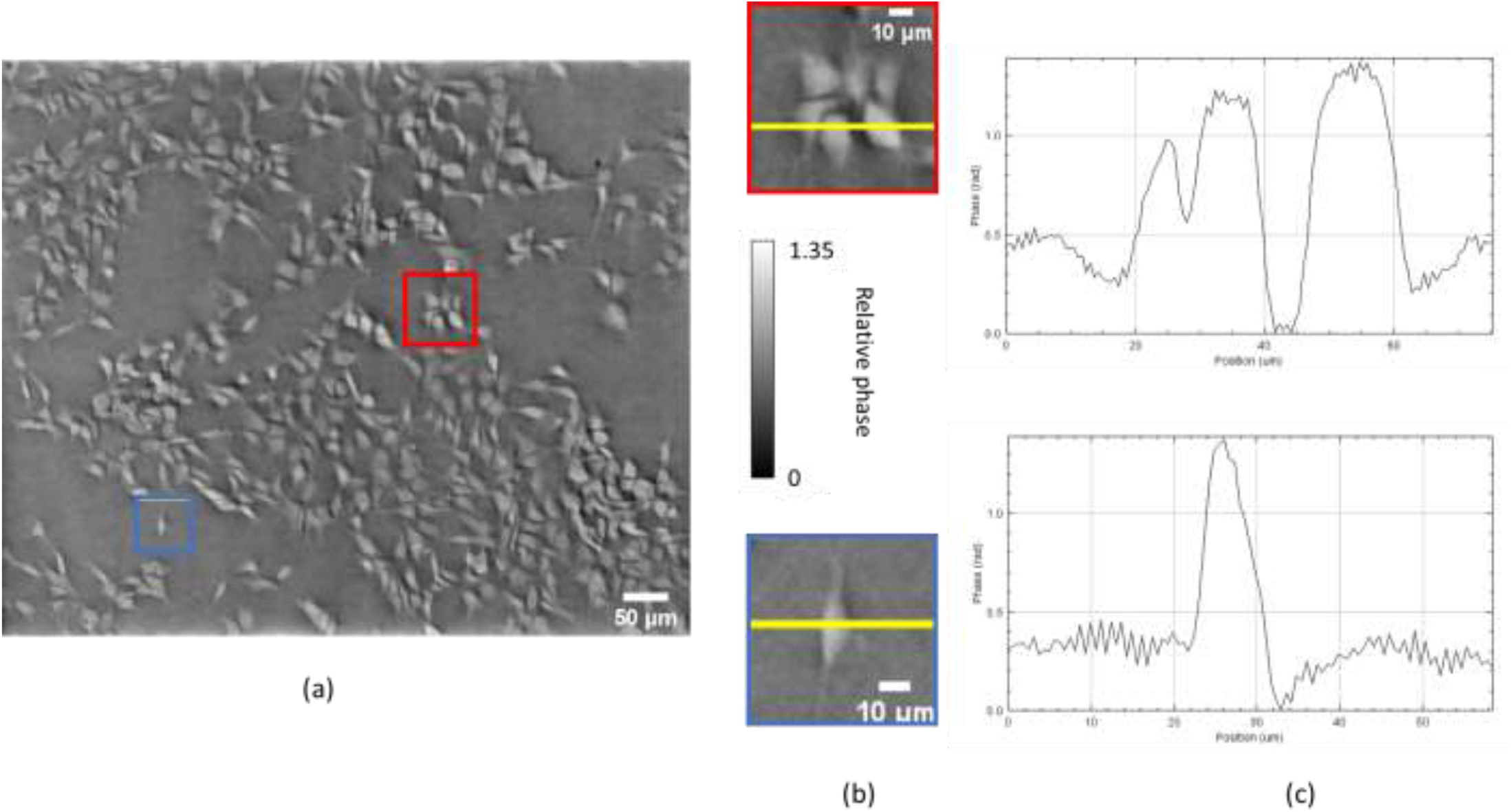
(a) pDPC phase contrast image of HEK cells images acquired using a 10x 0.3 NA objective lens with 0.36 NA illumination at 560 nm from an LED; (b): expanded images of the regions of interest marked in red and blue on (a); (c) phase profiles corresponding to yellow lines on (b).

Figure 4 shows a raw intensity image and the corresponding pDPC phase contrast image of a live *Caenorhabditis elegans* adult acquired using a 0.4 NA 20x microscope objective lens. The pDPC image provides superior contrast, clearly resolving single cells. The capability to study dynamics using pDPC is illustrated in Supplementary movie 1, which shows a 2.4 second clip of a live *C. elegans* acquired at a frame rate of 63 frames/s using the same objective lens. The arrow indicates the approximate position of the grinder, which is seen to move during pharyngeal pumping as the animal feeds.

**Figure 4.**
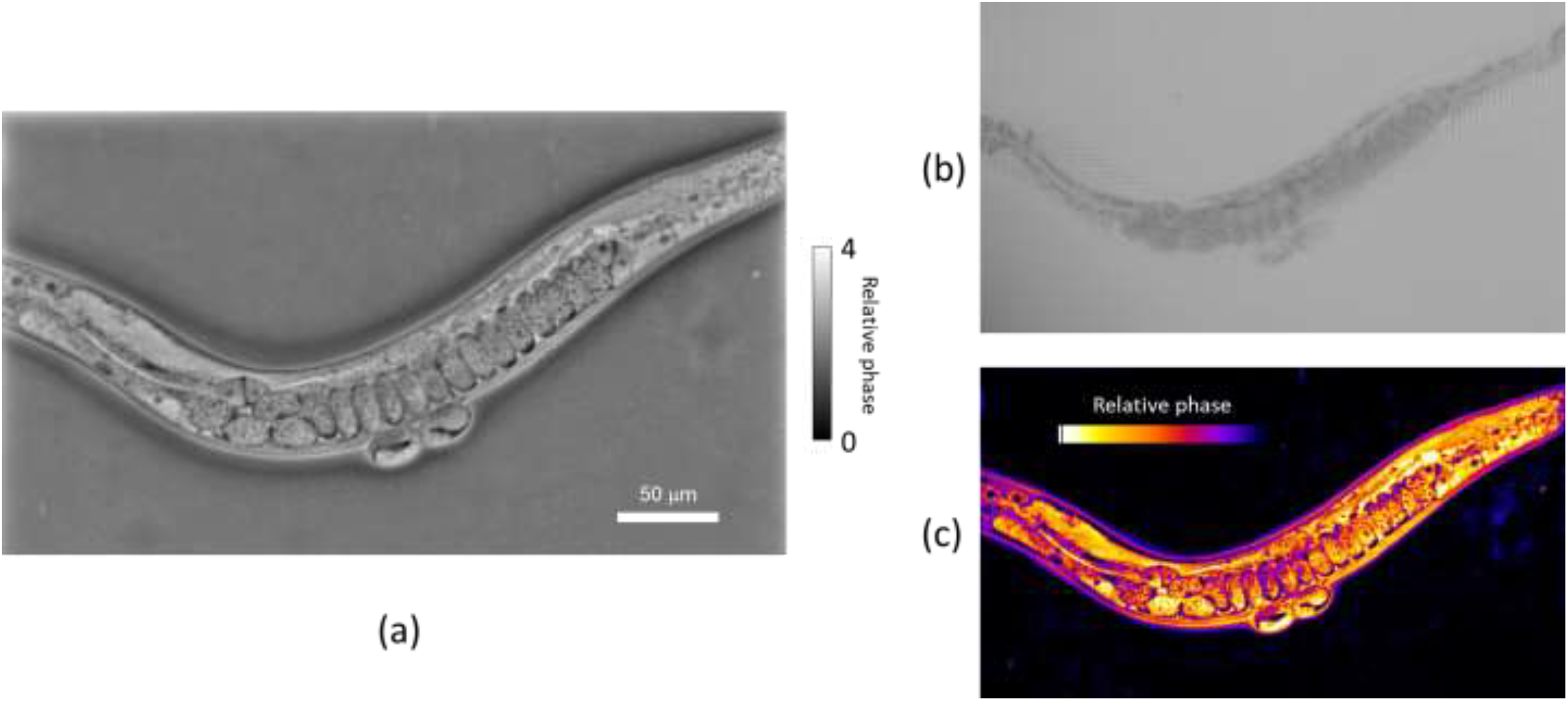
(a) pDPC image and (b) raw transmitted light intensity image of live C. elegans adult. (c) shows a 3D contour plot of the data presented in (a). This image was acquired with a 20x 0.4 NA objective lens with 0.36 NA illumination.

Figure 5 illustrates the capability to acquire co-registered phase contrast and fluorescence images of the same field of view applied to a live transgenic *C. elegans* adult in which epidermal stem cell nuclei are labelled with GFP expressed in the lateral seam cells. The same white light LED was used for transillumination and excitation. For the fluorescence image acquisition, an excitation filter (469 ±35 nm) was placed before the sample and the GFP fluorescence was detected above 520 nm.

**Figure 5.**
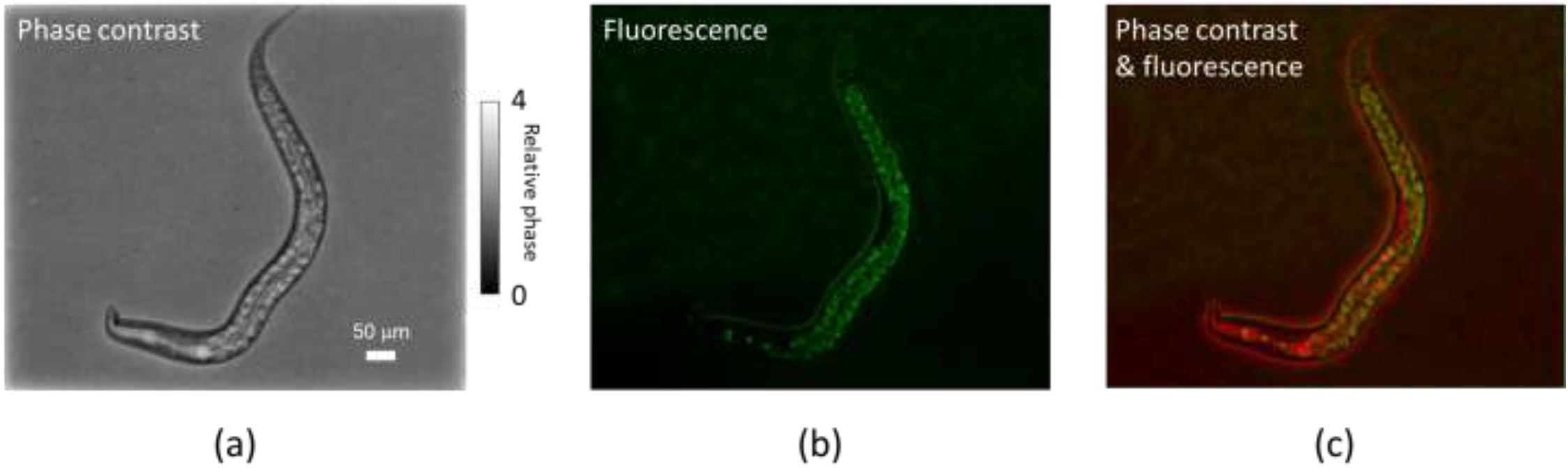
(a) pDPC phase contrast image, (b) GFP fluorescence image and (c) overlay of phase contrast (displayed in red) and fluorescence (green) images of live C. elegans adult images at x20 magnification. In panel b, seam cell nuclei can be seen in the head region while intestinal autofluorescence is seen in the rest of the animal body.

## Conclusions

In conclusion we have demonstrated a single-shot DPC phase contrast technique that utilises a polarisation-resolved camera sensor and can be implemented on most microscopes without requiring any specialized components beyond a custom polarisation mask to be located in the pupil plane of the condenser lens. This mask can be fabricated using low-cost polarizing film. The spectral versatility of this pDPC approach enables it to be implemented with multispectral imaging and it can be simply combined with fluorescence imaging, including to provide co-registered phase contrast and fluoresce images. The single shot pDPC image data acquisition can run at the maximum frame rate of the camera (75 frames/s) enabling phase contrast imaging of dynamic samples including live mobile organisms.

## Acknowledgements

The authors acknowledge helpful discussions with Lei Tian and Laura Waller. Significant components of the instrument reported here were co-designed and fabricated by Simon Johnson, Martin Kehoe nd John Murphy in the Optics instrumentation facility of the Physics Department at Imperial College London. We gratefully acknowledge funding from Cancer Research UK (A28450, A29368), Research England GCRF Institutional Award and the Imperial College London Impact Acceleration Accounts supported by the Biological Sciences Research Council (BBSRC EP/R511547/1). SK and YA are supported by funding from the Francis Crick Institute, which receives its core funding from Cancer Research UK (FC001003), the UK Medical Research Council (FC001003) and the Wellcome Trust (FC001003). Jonathan Lightley acknowledges a PhD studentship from the Engineering and Physical Sciences Research Council.

