## Supplementary figures and images for "Single-shot phase contrast microscopy using polarisation-resolved differential phase contrast"

### Supplementary movie 1

#### Slide 1
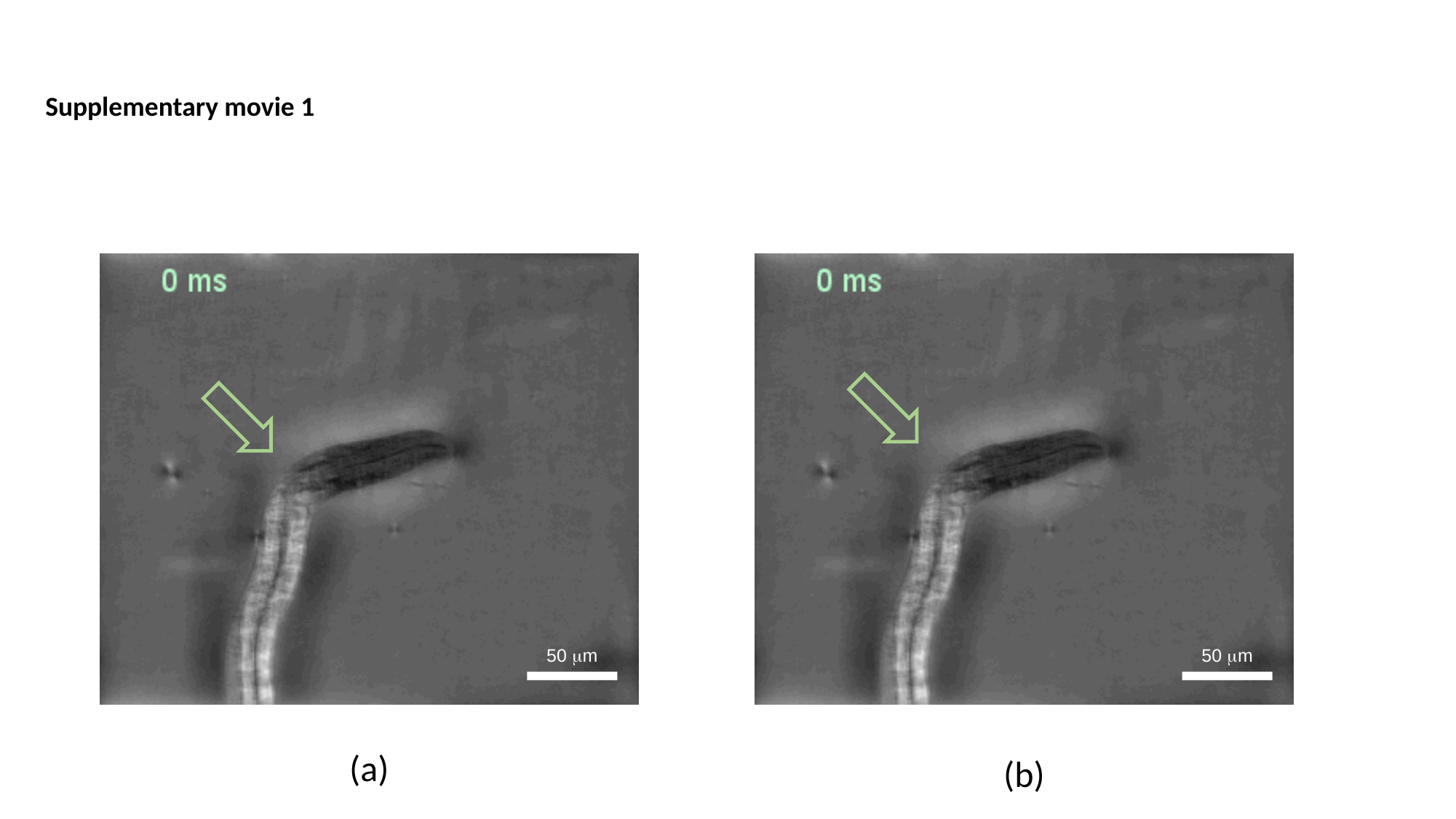

Supplementary movie 1
50 μm
50 μm
(a)
(b)
